# Polycomb Repressive Complexes occupancy reveals PRC2-independent PRC1 critical role in the control of limb development

**DOI:** 10.1101/2021.10.28.466236

**Authors:** Fanny Guerard-Millet, Claudia Gentile, Racheal Paul, Alexandre Mayran, Marie Kmita

**Affiliations:** Genetics and Development Research Unit, Institut de Recherches Cliniques de Montréal, Montréal, Québec, Canada; Department of Experimental Medicine, McGill University, Montréal, Québec, Canada; Molecular Genetics Research Unit, Institut de Recherches Cliniques de Montréal, Montréal, Québec, Canada; Department of Medicine, Université de Montréal, Montreal, Quebec, Canada

## Abstract

The Polycomb Repressive Complexes (PRC) are key players in the regulation of tissue-specific gene expression through their ability to epigenetically silence developmental genes. They are subdivided into two multicomponent complexes, PRC1 and PRC2, functioning through post-translational modifications of histone tails. A large body of work revealed functional interactions between PRC1 and PRC2, whereby trimethylation of lysine 27 on histone H3 (H3K27me3) by PRC2 contributes to the recruitment of canonical PRC1 (cPRC1). In parallel, a PRC2-independent binding of PRC1 has been uncovered and referred to as non-canonical PRC1 or variant PRC1 (vPRC1), in which PRC1-dependent ubiquitination of lysine 119 on histone H2A is involved in recruiting PRC2/propagating PRC2-dependent H3K27 trimethylation. While it was initially assumed that cPRC1 and vPRC1 bind distinct targets, subsequent evidence pointed to cPRC1 and vPRC1 sharing a significant subset of targets. How the functional interplay between PRC2, cPRC1 and vPRC1 contributes to gene regulation remains partially understood. Here, we show that, in the developing limb, PRC2 inactivation barely affects PRC1 occupancy, as the majority of PRC2- bound loci are bound by vPRC1 (RYBP-PRC1), both in wild type and PRC2 mutant limbs. Consistent with this, we found that loci bound by CBX2, a PRC1 subunit involved in the recognition of H3K27me3 and thereby recruitment of cPRC1, are, for the vast majority, also bound by vPRC1. Intriguingly, analysis of PRC2 mutant limbs revealed that while a large part of CBX2 occupancy is lost in absence of PRC2 function, as expected from the absence of H3K27me3, there is a significant number of genes retaining CBX2 occupancy as well as a few genes with apparent gain of CBX2 binding. Importantly, among these genes, 56 of them correspond to developmental genes known for playing a key role in limb morphogenesis. Based on the importance of vPRC1 in gene silencing, our findings emphasize the primary role of PRC2-independent PCR1 function in regulating developmental genes and questions the role of PRC2/cPRC1 in controlling developmental programs.

## Introduction

Polycomb group (PcG) proteins are known for their contribution to the silencing of developmental genes and thereby regulation of cell-fate decisions. PcGs were initially discovered in *Drosophila* for their role in repressing the *Hox* genes, which in turn regulate proper body segmentation during embryonic development (Jurgens, 1985; Lewis, 1978; Struhl, 1981). The PcG proteins form two functionally distinct multisubunit protein complexes, Polycomb Repressive Complex 1 and 2 (PRC1 and PRC2, respectively), each known for their inherent catalytic activities. PRC1 catalyzes the monoubiquitination of lysine 119 on histone H2A (H2AK119ub1) via the E3 ligase activity of RING1A/B, whereas PRC2 contains EZH1/2 which confers a methyltransferase activity responsible for the trimethylation of lysine 27 on histone H3 (H3K27me3) (Simon and Kingston, 2013). In vertebrates, the core components of PRC2 consist of Enhancer of Zeste Homologue 1 or 2 (Ezh1/2), Embryonic Ectoderm Development (Eed), and Suppressor of Zeste 12 (SUZ12). The PRC1 complex on the other hand, is characterized by extensive protein diversity and can be sub-classified into canonical and non-canonical/variant PRC1 (cPRC1 and vPRC1, respectively). The core of both cPRC1 and vPRC1 complexes consists of RING1A/B and a mutually distinct Pcgf protein (PCGF1-6). cPRC1 contains a CBX subunit (CBX2/4/6/7 or 8) as well as PCGF2 or PCGF4 and a PHC protein (PHC1/2/3) (Aranda et al., 2015). In contrast, vPRC1 does not contain a CBX protein and instead comprises RYBP or YAF2 and assembles into distinct complexes by associating with PCFG1/3/5/6 (Gao et al., 2012; Morey et al., 2013; Tavares et al., 2012). The presence of additional protein subunits results in diverse forms of PRC1 (cPRC1 and vPRC1) as well as PRC2, which may contribute to modulating their activity and function (reviewed in Blackledge and Klose, 2021; Schuettengruber et al., 2017).

In light of the diversity that exists amongst PRC complexes, great efforts have been made to elucidate how specific PRCs function in gene silencing to govern cell fate decisions. A widely accepted hierarchical recruitment model postulates that PRC2 once targeted, deposits H3K27me3 which is read by the CBX subunit of cPRC1, resulting in H2AK119ub deposition (Cao et al., 2002; de Napoles et al.; Min et al.; Wang et al.). This model predicts a direct communication between PRC2 and PRC1 leading to PRC1-mediated chromatin compaction and Polycomb domain formation, a function deemed critical for gene repression (Boyer et al.; Eskeland et al.; Francis et al.; Kundu et al.; Oksuz et al.; Schoenfelder et al.). Importantly, PRC1 can also be recruited independently of PRC2, further emphasized by the elucidation of vPRC1 (Almeida et al.; Blackledge et al.; Farcas et al.; Gao et al.; Schoeftner et al.; Tavares et al.). Moreover, PRC2 subunits can recognize and bind the H2AK119ub mark catalyzed by PRC1 allowing the spreading of PRC2-dependent H3K27me3 histone modification (Cooper et al.; Kalb et al.; Kasinath et al.). This in turn, has illuminated a predominant role for vPRC1 and H2AK119ub in gene repression (Fursova et al.). As continuous efforts are pursuing the functional roles of PRCs in various developmental contexts, it is becoming increasingly clear that PRC1 and PRC2 targeting mechanisms likely rely on histone modifications that can create feedback mechanisms that govern Polycomb domain formation and gene regulation (Blackledge et al.; Perino et al.; Rose et al.; Scelfo et al.).

Based on the distinct modes of cPRC1 and vPRC1 recruitment, it was initially assumed that they have different sets of targets, at least in a given cell population. However, with the emergence of better quality ChIP-seq assays, evidence was obtained for a strong overlap in sites occupied by cPRC1 and vPRC1 (Fursova et al.; Scelfo et al.; Zepeda-Martinez et al.), raising the possibility that a cPRC1/vPRC1 interplay contributes to PRC-dependent transcriptional control. Here, by studying PRC2 inactivation in the developing limb, we found that the binding of RING1B, the PRC1 catalytic subunit most expressed in the developing limb, is barely altered by PRC2 loss of function and PRC1 function, as evaluated by H2AK119ub genome-wide occupancy, remains largely unaffected. Moreover, in the wild type context, most PRC2-bound loci are occupied by RYBP, a vPRC1-specific subunit, suggesting that, in the developing limbs, the same loci can either be bound by cPRC1 or vPRC1, reminiscent of the situation previously identified in ES cells (Fursova et al.; Scelfo et al.). Accordingly, comparison of the genome-wide occupancy of CBX2, a cPRC1-specific subunit, and of RYBP, reveals that most of CBX2-bound promoters are also bound by RYBP in wild type limb buds. Further analysis of CBX2 occupancy in PRC2 mutant limb buds, lacking H3K27me3, reveals that while most CBX2 binding is lost, as expected from the absence of H3K27me3, there is a significant number of loci retaining CBX2 occupancy and few loci gaining CBX2 binding. Importantly, these loci retaining CBX2 occupancy comprise genes with demonstrated function in limb morphogenesis, including *Shh* and *Fgf8,* the genes encoding the two main signaling molecules underlying limb development (Zeller et al., 2009), while loci with gained CBX2 binding include the subset of *HoxA* genes required for limb patterning (Zakany and Duboule, 2007). Together, our data uncover the key role of PRC1 in controlling the limb developmental program, which can largely function in a PRC2-independent manner.

## Results

### PRC1 and PRC2 binding in the developing limb establishes distinct promoter categories genome-wide

To gain insights into PRC1 and PRC2 binding during limb development, ChIP-seq experiments for RING1B (PRC1) and SUZ12 (PRC2) were performed in proximal and distal limb buds dissected at embryonic day 12.5 (E12.5) (Fig. 1A). Genome-wide ChIP-seq analysis of PRC1/2 signal intensities around transcriptional start sites (TSSs +/- 1.5kb, referred to as promoters hereafter) in proximal and distal limbs, lead to the observation that more than half of promoters are bound by PRC1 and/or PRC2 (referred to as PRC-bound promoters), with 59% in proximal limb and 56% in distal limb (Fig. 1B). Further analysis revealed a partitioning of these PRC-bound promoters into 3 distinct categories. The first category comprises promoters occupied by SUZ12 and RING1B, category 2 consists of few promoters occupied by SUZ12 alone while promoters occupied by RING1B alone form category 3 (Fig. 1C). Of note, category 2 promoters correspong to loci with rather small SUZ12 ChIP peaks and we cannot exclude that, due to technical limitations, RING1B binding is not detectable. The proportions of promoters making up each category are similar in the proximal and distal limb and the largest proportion of PRC-bound promoters (79%) are exclusively occupied by PRC1 (category 3; Fig. 1C).

**Figure 1.**
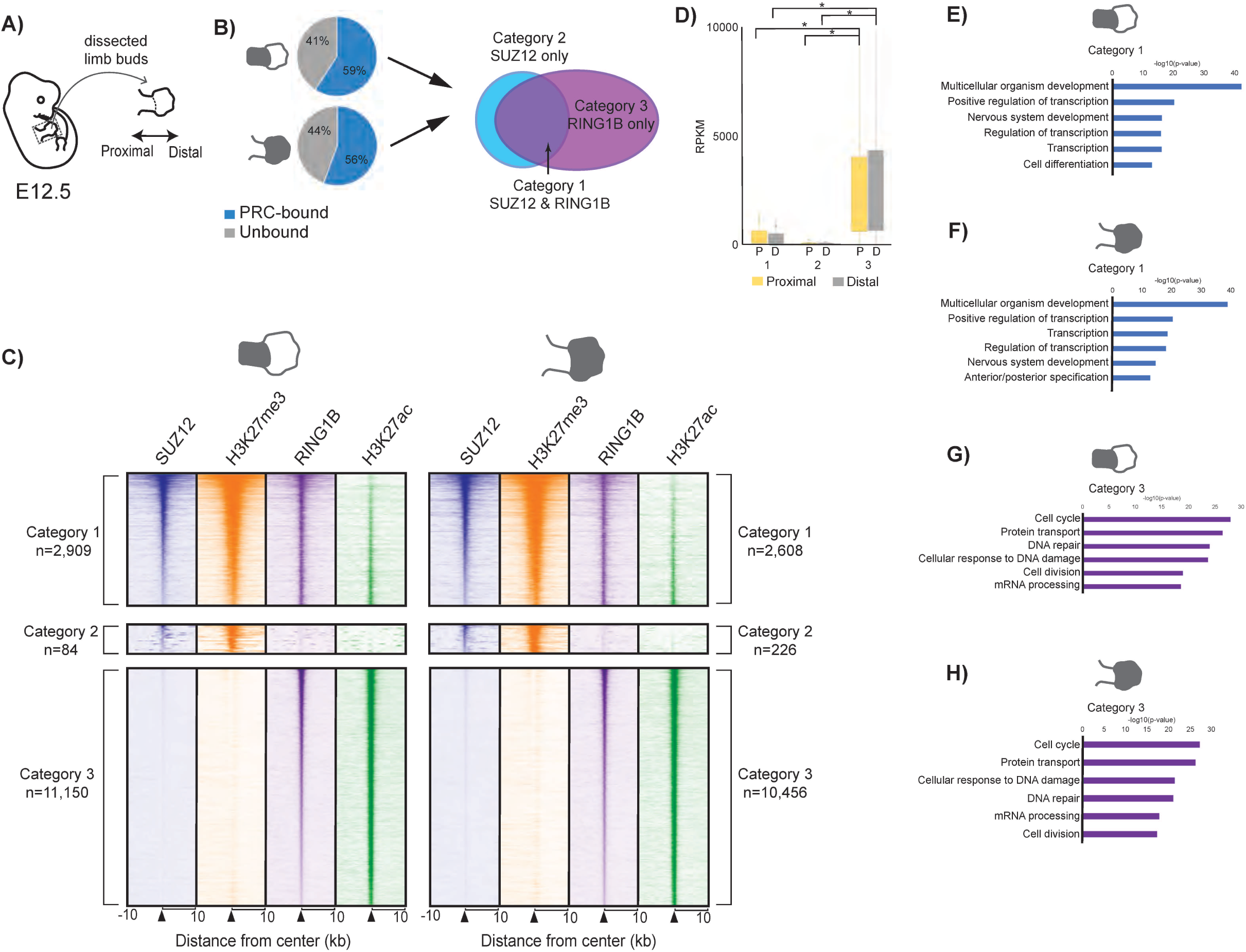
Genome-wide analysis of PRC occupancy in the developing limb reveals 3 distinct promoter categories. **(A)** Schematic of the dissection performed to isolate proximal and distal limb bud tissues. **(B)** Pie charts displaying the proportion of PRC-bound vs. unbound promoters genome-wide in proximal (upper) and distal (lower) limb tissues. Arrows from PRC-bound promoters point to a diagram representing how these promoters are further broken down into 3 categories according to PRC1 and PRC2 occupancy. To note, the grey shaded areas of the limb buds represent the corresponding tissues, that is either proximal or distal. **(C)** Heatmaps of ChIP-seq data showing the signal intensity of SUZ12 (blue), H3K27me3 (orange), RING1B (purple), and H3K27ac (green) at gene promoters, in the proximal and distal limb, for the 3 categories of promoters described. The number (n) of promoters for each category is indicated. **(D)** Box plot of gene expression (using RPKM values) from RNA-seq analysis in the proximal (P) and distal (D) limbs of each category of genes (1-3). Significant differences (p-value <0.05) in gene expression are depicted by an asterisk and were calculated using an unpaired t-test. **(E-H)** Gene Ontology terms (using DAVID) of top 6 biological processes associated with genes from categories 1 (blue bar graphs) and 3 (purple bar graphs) in the proximal and distal limb.

SUZ12 and RING1B occupancy at category 1 promoters correlates with the presence of the H3K27me3 repressive histone mark (Fig. 1C). Importantly, within this category, H3K27me3 as well as H3K27ac signals are detected at a subset of promoters, which reflects the cell heterogeneity of the limb tissues as these two histone marks are mutually exclusive (Pasini et al.; Tie et al.) (Fig. 1C). Category 1 promoters can thus be sub-divided into a sub-category referred to as fully-silenced promoters, that are bound by SUZ12, RING1B and are devoid of H3K27ac (i.e. 1459/2909 in proximal limb and 1525/2608 in distal limb) and a category of promoters reflecting a heterogeneous state: repressed in a subset of cells and active in the other subset (referred to as heterogeneously-silenced promoters hereafter) (Fig. S1A). Next, few promoters are exclusively occupied by SUZ12 in either limb tissue (category 2) and these are characterized by the presence of H3K27me3 and absence of H3K27ac (Fig. 1C). Category 3 promoters are characterized by strong RING1B and H3K27ac ChIP-seq signal intensities and are completely devoid of H3K27me3, consistent with the absence of PRC2 at these promoters (Fig. 1C). As such, category 3 promoters likely correspond to promoters occupied by the non-canonical/variant PRC1 complex (vPRC1), which does not depend on PRC2/H3K27me3 for its recruitment to DNA (Almeida et al., 2017; Blackledge et al., 2014; Tavares et al., 2012). To note, since the vast majority of RING1B- bound promoters belong to category 3 (79%), this suggests a high prevalence of vPRC1 occupancy at promoters in the developing limb.

RNA-seq analysis in wild type proximal and distal limb buds reveals that category 3 promoters coincide with highly expressed genes, in contrast to those from categories 1 and 2 (Fig. 1D). The high expression level for category 3 genes correlates with high H3K27ac signal intensity, while the low gene expression for category 1 and 2 correlates with the presence of H3K27me3. Amongst the genes belonging to category 1, those showing a higher RNA-seq signal correspond to the heterogeneously-silenced promoters in comparison to the fully-silenced promoters, which further emphasizes tissue heterogeneity (Fig. S1B). Furthermore, GO analysis revealed that promoters belonging to category 1 and 3 are enriched for distinct biological processes (Fig. 1E- H). Due to the small number of promoters in category 2, GO analysis did not reveal any significant enrichment for specific biological processes. A strong enrichment for biological process terms associated with development and transcriptional regulation are found for category 1 promoters (Fig. 1E and F). This is consistent with PRC function in silencing genes encoding developmental transcription factors (Boyer et al., 2006; Ku et al., 2008). In contrast, category 3 promoters, bound exclusively by RING1B, relate to genes involved predominantly in cell cycle, DNA repair and protein transport (Fig. 1G and H). These are terms that have been reported to associate with vPRC1 targets (van den Boom et al., 2016).

The observation that category 3 promoters, bound exclusively by RING1B, correlates so strikingly with high gene expression lead us to question whether promoters not bound by PRC (Fig. 1B) similarly correspond to highly expressed genes. Quite surprisingly, the unbound promoters demonstrated near-base line levels of gene expression when compared to category 3 gene expression (Fig. S1C-D). GO analysis for these unbound promoters represent biological process terms associated with sensory perception and signal transduction, amongst other similar top hits (Fig. S1E-F). Thus, these genes do not require PRC function at their promoters to be in a transcriptionally “silent” state (at least in the developing limb) while the most transcriptionally active genes genome-wide are genes for which we observe RING1B occupancy (category 3). Whether there is a functional link between vPRC1 and this high transcriptional activity (category 3) remains unclear as our data cannot establish if vPRC1 binding and acetylation of H3K27 occur in the same cells. Moreover, at least in ESCs, there is very little evidence that vPRC1 complexes contribute to active transcription (Fursova et al., 2019), though in some contexts this has been reported (Cohen et al., 2018; Gao et al., 2014; Morey et al., 2013; van den Boom et al., 2016). One possible explanation is that the robust expression of category 3 genes in a subset of limb cells requires vPRC1 function to secure gene silencing in the other limb cell populations where these genes shouldn’t be expressed. This is consistent with the recent demonstration of PRC2-independent mechanisms of gene silencing (Blackledge et al., 2020; Tamburri et al., 2020).

### PRC1 occupancy remains largely unaffected in PRC2 deficient limb buds

As PRC2-dependent trimethylation of H3K27 is recognized by the CBX subunits of PRC1, leading to cPRC1 recruitment (Cao et al.; Min et al.; Wang et al.), we wanted to establish to what extent PRC2 inactivation affects PRC1 occupancy. We thus decided to study embryos carrying the *Eed* conditional inactivation in the developing limb (referred to as *Eedc/-* hereafter; (Gentile et al.)) and first performed ChIP-seq for RING1B, the most highly expressed PRC1 catalytical subunit in wild type limbs at E12.5 (suppl. Fig. S2A). To verify that the *Eed* conditional inactivation in the developing limb efficiently results in PRC2 inactivation, we monitored H3K27me3 occupancy, which revealed that H3K27me3 is indeed lost in *Eed* mutant limbs (**Fig. 2A**). As H3K27me3 loss was observed in both the proximal and distal limb (Fig. S2B), we focused subsequent studies solely on the distal limb domain. Strikingly, we found that in the absence of PRC2 function, RING1B occupancy is barely altered in E12.5 distal limb buds (**Fig.2A**). We next verified if RING1B occupancy in PRC2 mutant limb coincides with a functionally active PRC1 by analyzing H2A ubiquitination at Lysine 119 (H2AK119Ub), the post-translational modification associated with PRC1 function (de Napoles et al.; Wang et al.). ChIP-seq for H2AK119ub both in wild type and *Eed* mutant limbs, reveals a highly similar H2AK119ub pattern, albeit a slightly overall reduced signal in *Eed* mutant limbs (**Fig.2A**). As H2AK119ub is associated with PRC1 repressive function (Blackledge and Klose, 2021) these results suggest that there is maintenance of PRC1-dependent gene silencing in PRC2 mutant limb. Consistent with this, while analysis of RNA-seq data in both wild type and *Eedc/-* E12.5 limb buds reveals a subset of deregulated genes (**Fig.2B**), overall changes in gene expression upon PRC2 inactivation affect a rather limited set of genes considering the number of gene promoters occupied by PRC2 in the wild type limb (**Fig.2B**).

**Figure 2.**
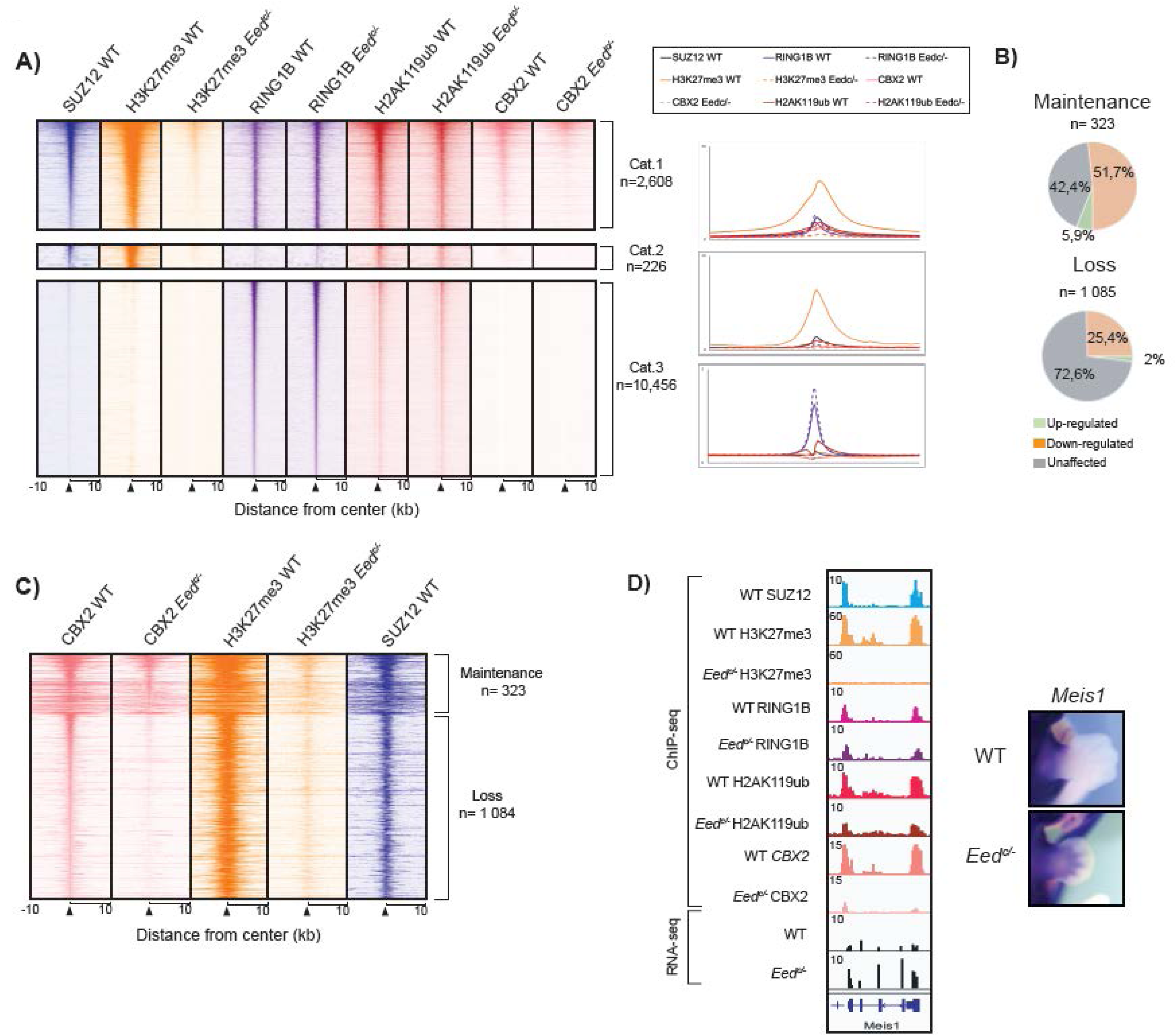
PRC1 function is largely unaffected by PRC2 inactivation in the developing limb. **(A)** Heatmaps of ChIP-seq data for SUZ12 (blue), H3K27me3 (orange), RING1B (purple), H2AK119ub (dark red) and CBX2 (pink) in wild type and *Eed* conditional mutant distal limb buds at E12.5 (left panel). Data are presented according to the 3 promoter categories identified (Fig.1). Average profiles of ChIP-seq data are presented on the right panel. **(B)** Pie charts summarizing the RNA-seq data at genes with maintained CBX2 binding (upper panel) and genes with loss of CBX2 occupancy (lower panel) upon PRC2 inactivation. Note the low number of upregulated genes in the two categories (5.9% and 2%, respectively). **(C)** Heatmaps of ChIP-seq data for CBX2 (pink), H3K27me3 (orange) in both wild type and *Eedc/-* distal limb buds at E12.5 as well as for SUZ12 (purple) in the wild type context, aligned according to maintenance of CBX2 occupancy in *Eedc/-* distal limb. Note that the intensity of CBX2 ChIP- seq peaks in *Eedc/-* tissue at promoters maintaining CBX2 binding appears slightly weaker that in the wild type distal limb but the significance of this small difference is difficult to assess as the ChIP-seq assays used are not quantitative. **(D)** Screenshots of IGV tracks at *Meis1*. Note the complete loss of H3K27me3, almost complete loss of CBX2 occupancy and gain of *Meis1* expression upon *Eed* inactivation (left panel). Whole- mount *in situ* hybridization for *Meis1* in both wild type and *Eed* mutant limbs are shown (right panel)

In addition, we analyzed CBX2 genome-wide binding as a proxy for cPRC1 occupancy in both wild type and *Eed* mutant limbs. As expected, CBX2 ChIP-seq data show a clear loss of CBX2 binding upon PRC2 inactivation (**Fig.2A, 2C**). As expected from loss of PRC2 function, a subset of gene promoters deprived of CBX2 and PRC2 become upregulated in the developing limb as exemplified with *Meis2* gain of expression (**Fig 2D**). However, only a small subset of genes lacking CBX2 binding following PRC2 inactivation, are actually up-regulated (2%; **Fig.2B**). We also found a significant number of genes with maintained CBX2 binding (n=323, **Fig. 2B-C**) despite loss of H3K27me3 (**Fig.3A-C**), consistent with previous finidings suggesting the existence of a PRC2-independent targeting of CBX2 (Zhen et al.; Zhen et al.).

**Figure 3.**
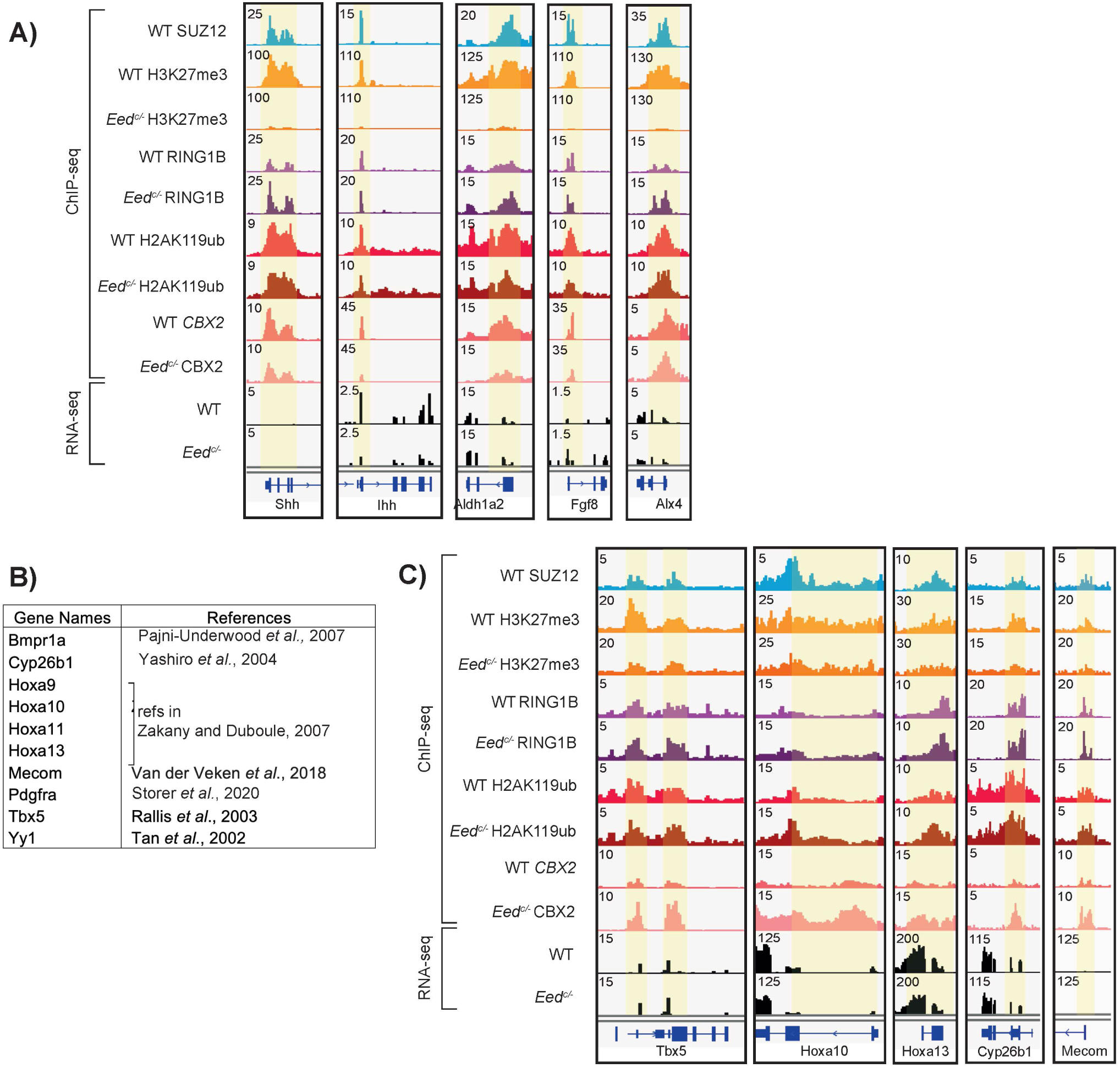
RYBP (vPRC1) occupancy is observed at PRC2 targets in the developing limb. **(A)** Screenshots of IGV tracks at a subset of representative genes showing maintenance of CBX2 occupancy in the *Eedc/-* limb bud. Note the unaffected RING1B occupancy despite loss of H3K27me3 and unaffected H2AK119ub occupancy as well as the absence of significant changes in expression levels. **(B)** List of the few genes showing an apparent gain of CBX2 binding upon PRC2 inactivation. Note that all these genes have demonstrated key roles in limb development, except *Yy1*, which has only a suspected role. **(C)** Screenshots of IGV tracks at a subset of genes showing an apparent gain of CBX2 occupancy despite PRC2 inactivation. Note the similar H2AK119ub profil in wild type and *Eedc/-* distal limb buds and identical expression levels.

### Gene promoters retaining CBX2 binding in PRC2 mutant limbs include key limb developmental genes

In an attempt to gain insights into the functional significance of genes retaining CBX2 binding in PRC2 mutant limbs, we analyzed the gene list in detail. Interestingly, we found that a large number of these genes are implicated in limb morphogenesis (**Table 1, Suppl. Table 1.**). These include *Sonic Hedgehog* (*Shh*) and *Fibroblast growth factor 8* (*Fgf8*), the signaling molecules underlying the function of the two main signaling centers in the developing limb, namely the zone of poralizing activity (ZPA) and the Apical Ectodermal Ridge (AER) (e.g. Benazet and Zeller, 2009). Overall, we found that among the genes retaining CBX2 binding in the PRC2 mutant limb (n=323), 46 genes were previously shown to be implicated in limb morphogenesis, formation of the limb vasculature or limb innervation (**Table 1).** IGV screen shots for a subset of these genes are shown in **Fig. 3A.** Of note, if anything, these genes are less expressed that in the wild type context and most show no significant change in expression level (**Fig. 3A**). For the other genes maintaining CBX2 binding in absence of PRC2, their potential role in limb development has not been investigated to our knowledge. Further studies will be necessary to establish if they, as we expect, contribute to proper limb formation.

**Table 1.** List of genes with demonstrated function in limb morphogenesis maintaining CBX2 binding in PRC2 mutant limb bud. References are provided as supplementary material.

| Gene Name | Reference | Gene Name | Reference |
| --- | --- | --- | --- |
| Aldh1a2 | {Niederreither et al., 2002} | Lmx1b | {Dreyer et al., 1998} |
| Alx1 | {Capellini et al., 2010} | Meis1 | {Mercader et al., 1999} |
| Alx3 | {Matsumoto et al., 2013} | Pax3 | {Bober et al., 1994} |
| Alx4 |  | Lhx9 | {Tzchori et al., 2009} |
| Atoh1 | {Yuengert et al., 2015} | Pax9 | {Peters et al., 1998} |
| Barx1 | {Church et al., 2005} | Pitx1 | {Szeto et al., 1999} |
| Dlx6 | {Robledo et al., 2002} | Prrx1 | {Logan et al., 2002} |
| Ebf1 | {El-Magd et al., 2021} | Ptf1a | {Orchard et al., 2019} |
| Ecel1 | {Nagata et al., 2017}; {Jin et al., 2020} | Runx1 | {Wang et al., 2005} |
| En1 | {Wurst et al., 1994} | Runx2 | {Yoshida et al., 2004} |
| En2 | {Davis et al., 1991}; {Huettl et al., 2015} | Sall3 | {Kawakami et al., 2009} |
| Fgf4 | {Niswander and Martin, 1993} | Shh | {Niswander et al., 1994} |
| Fgf5 | {Clase et al., 2000} | Six2 | {Yamamoto-Shiraishi and Kuroiwa, 2013} |
| Fgf8 | {Mahmood et al., 1995} | Sp6 | {Hertveldt et al., 2008} |
| Fgf9 | {Hung et al., 2007} | Sp8 | {Bell et al., 2003} |
| Flt1 | {Eshkar-Oren et al., 2009} | Sp9 | {Kawakami et al., 2004} |
| Foxc1 | {Fontana et al., 2017} | Tbx1 | {Dastjerdi et al., 2007} |
| Gdnf | {Wang et al., 2002} | Tfap2a | {Leslie et al., 2013} |
| Hand1 | {Firulli et al., 2017} | Tfap2b | {Satoda et al., 2000} |
| Hic1 | {Carter et al., 2000} | Tfap2c | {Tsujimura et al., 2015} |
| Hoxa11os | {Kherdjemil et al., 2016} | Twist1 | {Hirsch et al., 2018} |
| Ihh | {Vortkamp et al., 1996} | Wnt10b | {Ugur and Tolun, 2008} |
| Lhx2 | {Tzchori et al., 2009} | Zic3 | {Quinn et al., 2012} |

Unexpectedly, we also found few genes that gain CBX2 binding in PRC2 mutant limbs (**Fig. 3B-C**). These include genes from the *HoxA* cluster that are known for their role in limb patterning, namely *Hoxa9, Hoxa10, Hoxa11* and *Hoxa13* (e.g. Zakany and Duboule) as well as *Tbx5* whose loss of function results in complete absence of forelimbs (Rallis et al.) and *MECOM,* whose mutation in humans results in limb deformities (van der Veken et al.). It also includes *Yy1*, a transcription factor belonging to the GLI-Kruppel class of zinc finger proteins, whose function is suspected to be associated with limb development (Tan et al.), but has yet to be formally demonstrated. Together, these data indicate that PRC2-independent CBX2 occupancy is likely critical for proper limb development, but further studies will be necessary to fully understand the role of CBX2 in limb formation.

### PRC2 targets can be bound by vPRC1 in developing limbs

Based on the largely unaffected RING1B occupancy detected in PRC2 mutant limbs, we next examined whether the maintenance of RING1B binding is due to the ectopic recruitment of vPRC1. To address this point, we performed ChIP-seq for RYBP, a vPRC1-specific subunit (Gao et al.; Tavares et al.). Comparison between wild type and PRC2 mutant limbs, does not show any significant ectopic binding of vPRC1 as most, if not all, PRC2 targets are also occupied by RYBP in the wild type limb (**Fig.4A**) indicating that PRC2 targets can be bound by vPRC1. As the ChIP-seq data are an average of SUZ12 and RYBP binding in the heterogenous limb cell populations, these data cannot discriminate between co-binding of PRC2 and vPRC1 or binding at the same loci but in different cell populations. To further confirm RYBP binding at PRC2 targets, we compared CBX2 and RYBP genome-wide occupancy, which shows that CBX2 targets can indeed be bound by RYBP in the wild type developing limb (**Fig. 2B**). Together, these data indicate that, in the developing limb, a gene promoter can either be bound by cPRC1 or vPRC1, reminiscent to the situation observed in ES cells (Fursova et al.; Scelfo et al.). However, based on our finding that CBX2 can be recruited independently of the presence of the H3K27me3 mark (**Fig. 2B; Fig. 4B**), it is difficult to know which aspects of CBX2 binding truly corresponds to cPRC1, more so as CBX2 binding occurs at developmental genes for which we also found RYBP (vPRC1) binding (**Fig. 4B**). It is also worth noting that developmental genes with *bona fide* functions during limb development (*Tbx5, Hoxa10, Hoxa13, Cyp26b1, Bmpr1a, Pdgfra*) or suspected function (*Yy1*) showing robust SUZ12 (PRC2) occupancy (*Tbx5, Hoxa10* and *Hoxa13*) or moderate SUZ12 binding (*Cyp26b1* and *Bmpr1a*) are not up-regulated in PRC2 mutant limb (**Fig. 3**). Based on the maintained H2AK119ub genome-wide occupancy despite PRC2 inactivation (**Fig. 2A**), it appears that the PRC-dependent control of the limb developmental program is primarily achieved by the PRC2-independent function of PRC1.

**Figure 4.**
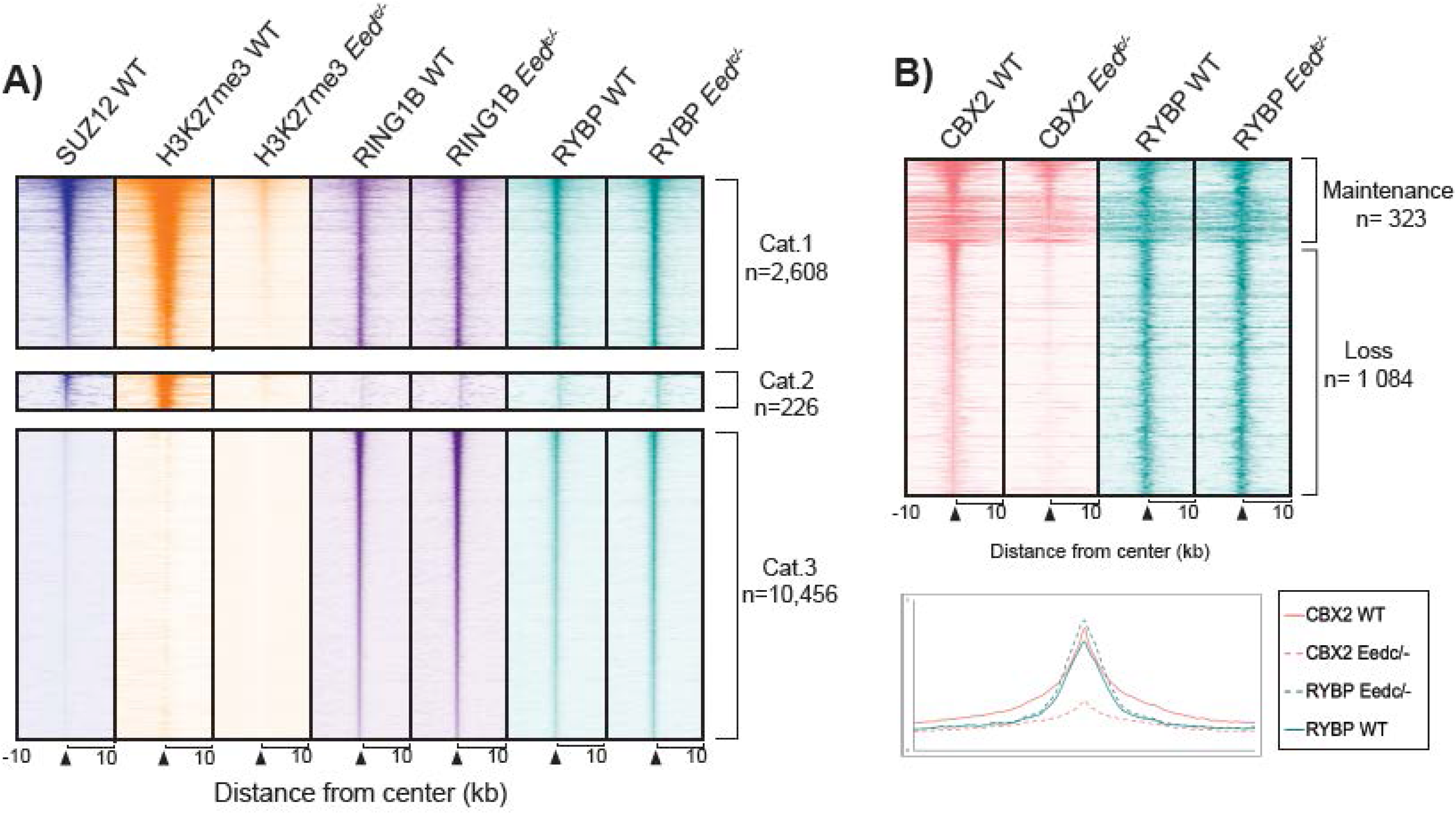
CBX2 occupancy is maintained despite PRC2 inactivation at developmental genes with essential functions for limb development. (A) Heatmaps of ChIP-seq data for SUZ12 (blue) in the wild type limb and H3K27me3 (orange), RING1B (purple) and RYBP) green) in wild type and *Eedc/-* distal limb buds, aligned according to the 3 promoter categories previously identified (Fig.1). Note RYBP (vPRC1) occupancy at all SUZ12 (PRC2) bound promoters and the identical RING1B and RYBP binding in both wild type and *Eedc/-* tissues. (B) Heatmaps of ChIP-seq data for CBX2 (pink) and RYBP (green) in the wild type and *Eedc/-* distal limb, aligned according to CBX2 ChIP-seq peak intensity. RYBP (vPRC1) occupancy is observed at all CBX2 (cPRC1) targets (upper panel panel). Average profiles of ChIP-seq data are shown below (bottom panel).

## Discussion

Polycomb Repressive Complexes (PRC) represent a critical family of epigenetic regulators of gene expression, notably during embryonic development. While PRC2 deposits the H3K27me3 repressive mark, PRC2-independent PRC1 occupancy silences genes by counteracting RNA polymerase II binding and transcription initiation (Blackledge and Klose). Based on the distinct mechanisms contributing to cPRC1 and vPRC1 recruitment, it was initially assumed that cPRC1 and vPRC1 have distinct sets of targets. Subsequent studies, however, provided evidence that in ES cells, there is a significant overlap between cPRC1 and vPRC1 targets (Fursova et al.; Scelfo et al.). Despite continuous significant advances in the field, the functional interplay between PRC1 and PRC2 remains partially understood.

Here, using the developing limb as an *in vivo* model system, we provide evidence that PRC2 inactivation barely affects PRC1 occupancy and function, as demonstrated by RING1B and H2AK119ub ChIP-seq data. Moreover, we found that, in the wild type context, vPRC1 binds cPRC1 targets, as defined by comparison between RYBP, PRC2 and CBX2 genome-wide occupancy. Although it remains unclear if vPRC1 and cPRC1 occupy PRC2 targets in the same cells, our results indicate that vPRC1 target genes include cPRC1 targets, at least for CBX2-containing cPRC1, in agreement with previous studies in ES cells (Fursova et al.; Scelfo et al.).

Consistent with the hierarchical recruitment of cPRC1 via CBX-dependent recognition of H3K27me3, we found that a large proportion of CBX2 genome-wide binding is lost in PRC2 mutant limbs. However, there is a significant number of promoters at which CBX2 binding is maintained in absence of any detectable H3K27me3 mark and even more strikingly, we uncovered ectopic CBX2 binding at a subset of developmental genes (**Fig. 3**). Together, these data reveal that CBX2-containing PRC1 doesn’t follow the canonical mechanism of cPRC1 recruitment. Analysis of the other CBX subunits, namely CBX4, 6, 7 and 8, in a PRC2 deficient context will be important to establish whether CBX2 is an atypical CBX subunit or whether the CBX subunits are not appropriate markers for the canonical recruitment. Importantly, the evidence that both CBX2-containing cPRC1 and vPRC1 share common targets and key limb developmental genes are not affected by PRC2 inactivation, raise the question of the functional significance of PRC2-dependent recruitment of PRC1. Further studies will be necessary to address this important question.

Our data shows that the majority of PRC target genes, in the wild type limb bud, are vPRC1-specific targets (category 3 genes). Moreover, PRC2/cPRC1 targets are also bound by vPRC1 and the vPRC1 binding at PRC2 targets most likely explains why we do not observe changes in H2AK119ub at most cPRC1 targets in PRC2 mutant limb buds. Yet, a subset of these target genes become up-regulated in PRC2 mutant limb buds, suggesting that PRC2 targets, which are primarily developmental genes, are silenced through distinct PRC-dependent mechanisms, some primarily requiring PRC2 function while vPRC1 function alone appears to be sufficient for silencing the other set of PRC2 targets. Of note, a significant number of genes that retain CBX2 binding in PRC2 mutant limb are genes previously demonstrated as having a critical function in limb development (**Fig. 3A**), while the function of the others has, to our knowledge, not yet being studied in the context of limb development. For genes with known function in limb formation, it is interesting to note that they expressed in specific sub-domains/sub-populations of cells of the developing limb, as exemplified with *Shh* and *Fgf8*. Moreover, for many of these genes, expression in inappropriate cells within the developing limb results in limb malformations as illustrated by *Shh* ectopic expression in the anterior limb margin leading to postaxial polydactyly (Lettice et al.). It is thus tempting to speculate that there is a safeguard mechanism provided by CBX2-containing PRC1 that compensates, at least partially, for the loss of PRC2 function at developmental genes whose functions are required in a specific subset of limb bud cells to ensure proper limb morphogenesis.

The apparent preponderant role of vPRC1 in developing limbs also raises the question of vPRC1 function at gene promoters associated with high transcription (category 3). Based on the robust ChIP-seq peaks for H2AK119ub and the evidence that non-PRC target genes show minimal or no transcriptional activity, we speculate that vPRC1 occupancy occurs at genes highly transcribed in a subset of limb cells, in which vPRC1 is not bound, while vPRC1 function ensures that these genes are not transcribed in cells where their expression could be detrimental for limb formation. Alternatively, some studies suggest that vPRC1 could be associated with transcriptional activation (Creppe et al.; Frangini et al.; Morey et al.) and we cannot fully exclude that part of PRC1 function in the developing limb is associated with transcriptional activation. However, we do not favor this hypothesis based on the unambiguous H2AK119ub occupancy at PRC1 target genes. Analyses of vPRC1, H2AK119ub and H3K27ac occupancy at single cell resolution will be required to discriminate between these two models.

As far as limb development is concerned, it is striking that *Rnf2* (encoding for RING1B, the main PRC1 catalytic subunit expressed in the developing limb) and *Eed* (PRC2) conditional inactivation results in a similar and rather mild phenotype (Gentile et al.; Wyngaarden et al.; Yakushiji-Kaminatsui et al.). In contrast, the combined inactivation of RING1A and RING1B abrogate almost completely limb formation (Yakushiji-Kaminatsui et al.), while *Ring1A* null mice develop almost normally (del Mar Lorente et al.). This suggests that RING1B is able to compensate for RING1A loss of function. It is worth mentioning that *Rnf2* (RING1B) and *Eed* conditional inactivation in developing limbs was achieved using *Prrx1:Cre*, a deletor transgene which results in full inactivation only at E11.5, i.e. two days after the initiation of limb development (Gentile et al.; Kmita et al.; Wyngaarden et al.; Yakushiji-Kaminatsui et al.). Thereby, the phenotypes associated with *Rnf2* and *Eed* conditional inactivations are, most likely, milder than what we could expect from the complete inactivation of these genes in nascent limb buds. A novel limb-specific Cre line, achieving complete gene inactivation at early limb bud stage is necessary to verify this hypothesis.

It has been shown that vPRC1 has more catalytic activity than cPRC1 and that cPRC1 actually contributes little to genome-wide Ub (Fursova et al.; Gao et al.; Rose et al.). These data supports a view whereby vPRC1 could have the capacity to substantially compensate for cPRC1/PRC2 inactivation. Yet, it is important to keep in mind that a subset of PRC2 targets become up-regulated upon *Eed* inactivation in the developing limb, suggesting that there might be different requirements for the silencing of developmental genes. Analysis of gene regulation in conditional compound mutants for both PRC1 and PRC2 will be required to gain further insights into the potential ability of vPRC1 to compensate, at least partially, for PRC2 loss of function, as was recently observed to occur to preserve epidermal stem cell identity (Cohen et al.).

One puzzling finding in this study is the evidence that CBX2 binding is actually gained at few limb developmental genes upon PRC2 inactivation (**Fig. 3B-C**). These include, for instance, the four *HoxA* genes contributing to limb patterning (Zakany and Duboule, 2007). How and why CBX2 occupancy at these genes becomes gained following PRC2 inactivation? It would be tempting to think that there is a possible developmental pressure to maintain the silencing of these genes in limb cells where they are normally not expressed. Indeed, ectopic expression of *Hoxa13* in the entire limb bud, a gene whose expression is normally restricted to the distal limb domain (presumptive digit domain), results in severe limb malformation, with an almost complete loss of the proximal limb skeleton (Williams et al., 2006). Yet, a similar phenotype was obtained for *Hoxd13* ectopic expression and our results show that not only there is no gain of CBX2 occupancy at *Hoxd13* in PRC2 mutant limb, it also looses CBX2 binding and becomes upregulated. Further studies are thus required to address the functional significance and mechanism(s) associated with the gain of CBX2 occupancy in absence of PRC2 function at the *HoxA* cluster in developing limbs.

Numerous studies in embryonic stem cells (ES cells) and *in vitro* ES cell differentiation systems have been key to understanding many aspects of PRC biology. Whether PRC functions similarly in other developmental systems remains however elusive. Our results reveal that many of the PRC features discovered in ES cells apply to the developing limb but also emphasize that PRC1- and PRC2-mediated gene regulation need to be studied taking into account the tight link existing between PRC1 and PRC2. Finally, our data point to an even more complex cross-talk underlying, at least in the developing limb, PRC1 and PRC2 recruitment mechanisms. In turn, it calls for a more precise definition of cPRC1. Either CBX2 cannot be considered as a cPRC1-specific subunit or cPRC1 can be recruited independently of PRC2. In this latter case, cPRC1 should not be considered as the PRC1 complex being recruited through the canonical mechanism of CBX recognition of the PRC2-dependent H3K27me3 histone mark.

## ACKNOWLEDGEMENTS

We thank Sarah Boissel and members of the molecular biology and functional genomics core facility at the IRCM for generating sequencing libraries. We are particularly grateful to Kmita lab members for helpful discussions and to Drs. Neil Blackledge, Raphaël Margueron and Nicole Francis for insightful comments on the manuscript and exciting discussion. Bioinformatics analyses were enabled in part by support provided by Calcul Quebec (www.calculquebec.ca) and Compute Canada (www.computecanada.ca). This work was supported by the Canadian Institute for Health Research (MOP-115127; MOP-174989) to MK. CG was supported by the Jacques Gauthier IRCM fellowship, AM was supported by the IRCM Challenge Fellowship and FGM is supported by the IRCM Challenge Fellowship.

## AUTHOR CONTRIBUTIONS

CG, FGM and MK conceived the study. CG and FGM designed and performed the experiments. AM performed the RNA-seq analysis. RP helped with the ChIP-seq assays. CG, FGM, AM, RP and MK analyzed and interpreted the data. MK wrote the manuscript with inputs from CG, FGM and RP.

## CONTACT FOR REAGENT AND RESOURCE SHARING

Further information and requests for resources and reagents should be directed to and will be fulfilled by the Lead Contact, Marie Kmita.

## EXPERIMENTAL MODEL AND SUBJECT DETAILS

The inactivation of *Eed* was generated using the limb specific *PrxCre* line (Logan et al., 2002) crossed with the previously generated *Eed* floxed mouse line (Yu et al., 2009) (Jax stock #022727).

## METHOD DETAILS

### Whole-mount in situ hybridizations

*In situ* hybridizations were carried out according to standard procedures as previously described (Gentile et al.). The RNA probe for Meis1 was as described previously (Mercader et al.). Briefly, embryos were rehydrated through a methanol series (100% - 30%), washed in PBST (0.1% Tween) and bleached for an hour on ice using 6% hydrogen peroxide before undergoing Proteinase K treatment for 15 minutes at RT. Next, embryos were hybridized with the RNA probes overnight at 68°C. Embryos were then washed and incubated with anti-DIG antibody (Cedarlane) overnight at 4°C. The coloration was done using BCIP and NBT (Roche). Embryos were genotyped prior to experimentation and WT and Eedc/- littermates were treated identically in the same assay for comparison. A minimum of 3 embryos per genotype was assayed for reproducibility (n=3).

### RNA Preparation and Sequencing

RNA was extracted from dissected proximal and distal limb buds from 3 independent pools of wild type limb buds (n=3) and from two independent pools of *Eed^c/-^* limb buds (n=2). The extraction was performed using the RNAeasy Plus mini kit. Next, mRNA enrichment, library preparation and flow-cell preparation for sequencing were performed by the IRCM Molecular Biology Core Facility according to Illumina’s recommendations. Sequencing was done on a HiSeq 2000 instrument with a paired-end 50 cycles protocol.

### Chromatin Immunoprecipitation (ChIP) and Sequencing

ChIP-seq experiments for SUZ12, RING1B, and H3K27ac were performed as previously described (Gentile et al.). ChIP-seq experiments for H3K27me3, H2AK119ub, CBX2 and RYBP in wild type and *Eed^c/-^* limb tissue were performed by crosslinking tissue with 1% formaldehyde for 13 minutes and sonicated to obtain fragments between 100-600 bp. Protein A and Protein G Dynabeads (Invitrogen) were incubated for 6 hours at 4°C with 5 μg H3K27me3 (07-449, Millipore), 5ug H2AK119ub (8240, Cell Signaling), 5ug CBX2 (ab80044, Abcam) and 5ug RYBP (ab3637, Millipore Sigma) and coupled to beads overnight at 4°C. The immunoprecipitated samples were then washed with different wash buffers depending on the antibody present. For H3K27me3 antibody, 3 sequential washes using LiCl (100mM Tris (pH 8), 500mM LiCl, 1% NP-40, 1% Na-Deoxycholate) were performed. For H2AK199ub, CXB2 and RYBP sequential washes in low salt (1% Triton, 0,1% SDS, 150 mM NaCl, 20 mM Tris (pH8), 2 mM EDTA), high salt (1% Triton, 0,1% SDS, 500 mM NaCl, 20 mM Tris pH8, 2 mM EDTA), LiCl (1% NP-40, 250 mM LiCl, 10 mM Tris (pH8), 1 mM EDTA) and TE buffer (50 mM NaCl, 10 mM Tris (pH8), 1 mM EDTA) were performed. The DNA was then purified on QIAquick columns (Qiagen). Library and flow cells were prepared by the IRCM Molecular Biology Core Facility according to Illumina’s recommendations and sequenced on Illumina Hiseq 2500 in a 50 cycles paired-end configuration.

### Quantification and statistical analysis

#### ChIP-seq Data Analysis

ChIP-seq reads were aligned to the mm10 genome using bowtie v.2.3.1 with the following settings: bowtie2 -p 8 --fr --no-mixed --no-unal –x. Sam files were converted into tag directories using HOMER v4.9.1 and into bam files using Samtools v1.4.1 view function. Analysis of genome-wide occupancy was done by identifying peaks for these datasets by comparing each sample to its input using MACS v2.1.1.20160309 callpeak function using the parameters: --bw 250 -g mm --mfold 10 30 -p 1e-5. Analysis of SUZ12, H3K27me3, RING1B and H3K27ac promoter occupancy was determined using signal intensities and by averaging values from ChIP-seq data replicates (for distal limb datasets of SUZ12 and RING1B only). Heatmaps and average profiles were generated using the Easeq software (http://easeq.net) (Lerdrup et al., 2016). ChIP-seq data were visualized on the IGV software using BigWig files generated using the makeUCSCfile HOMER command using the following parameters: -fsize 1e20 -fragLength 150 –bigWig. GO analysis was performed using the DAVID webtool (Huang da et al., 2009a, b).

#### RNA-seq Data Analysis

RNA-seq experiments were carried out as previously described (Gentile et al.). Paired-end reads were aligned to the mm10 reference genome using TopHat2 (Kim et al., 2013) with the parameters --rg-library “L” --rg-platform “ILLUMINA” --rg-platform-unit “X” --rg-id “run#many” --no-novel-juncs --library-type fr-firststrand -p 12. Gene expression was quantified with the HOMER analyzeRepeats command (Heinz et al., 2010) and differential expression was assessed with edgeR 3.12.1(McCarthy et al., 2012). MA plots were generated by averaging the wild type and *Eed^c/-^* counts per million (CPM) and plotting the log(CPM+1) against the log2 fold-change (log2FC) of wild type vs *Eed^c/-^* CPM values. The heatmap clustering and PCA was performed using the ClustVis webtool (Metsalu and Vilo, 2015).

---

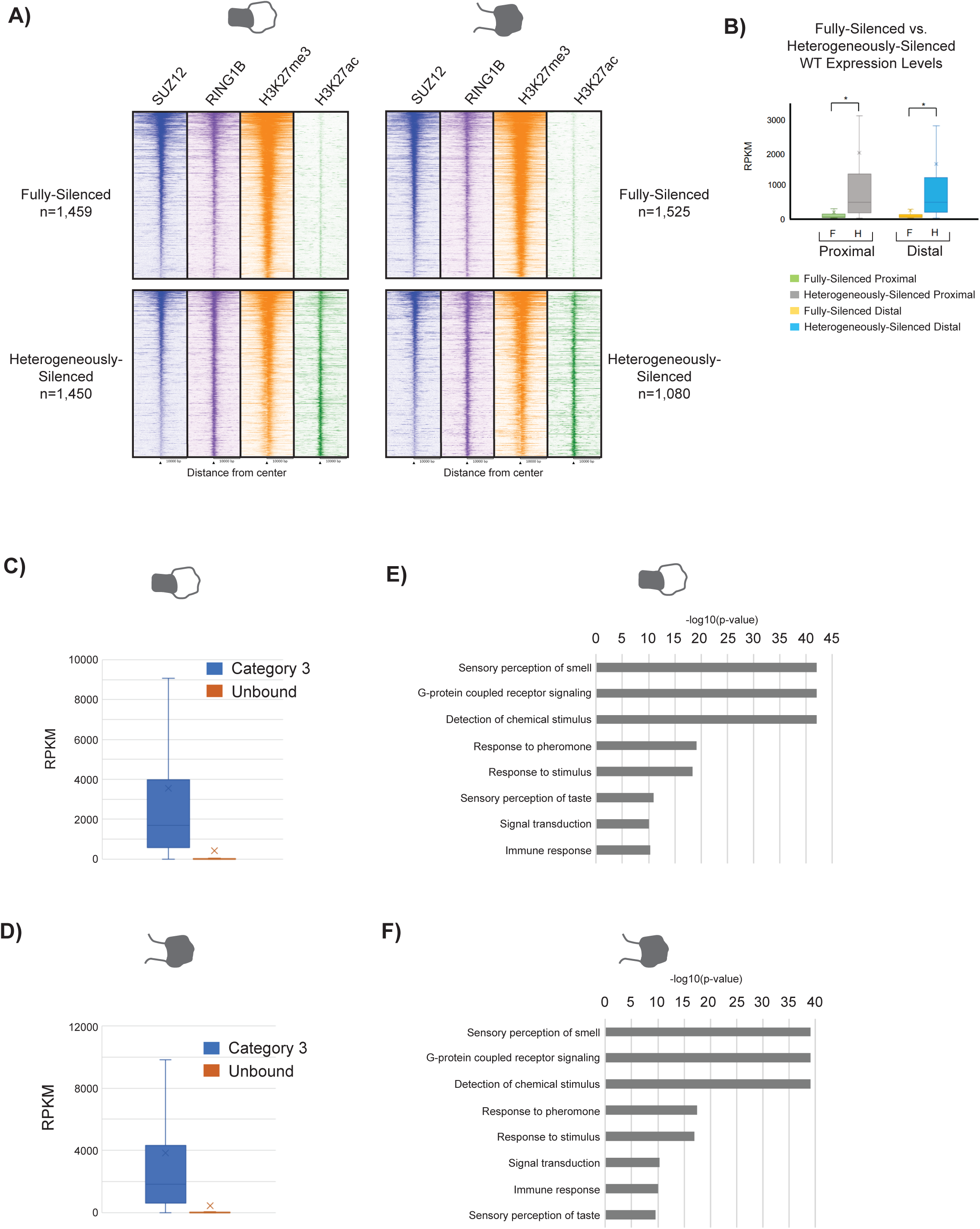

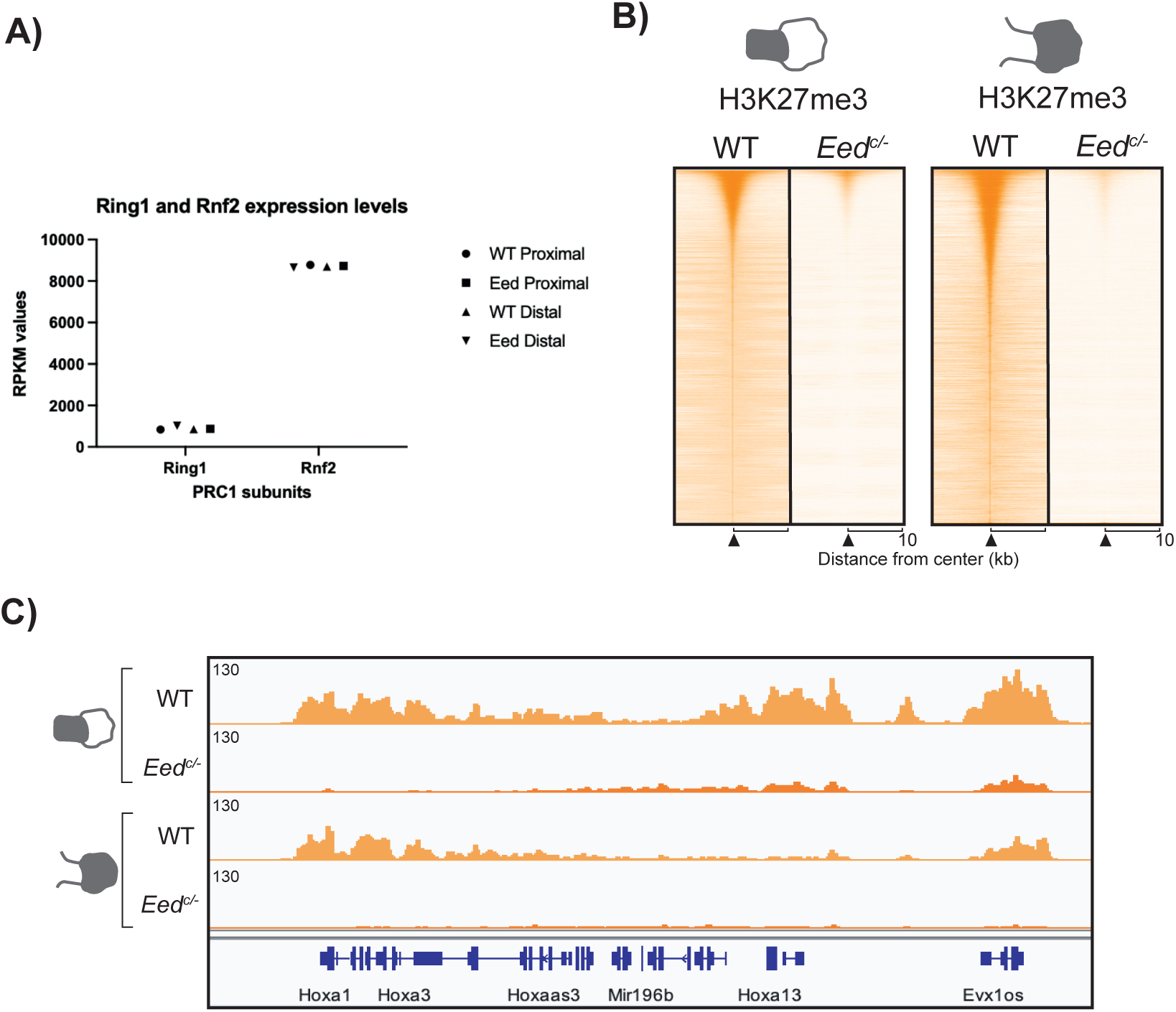

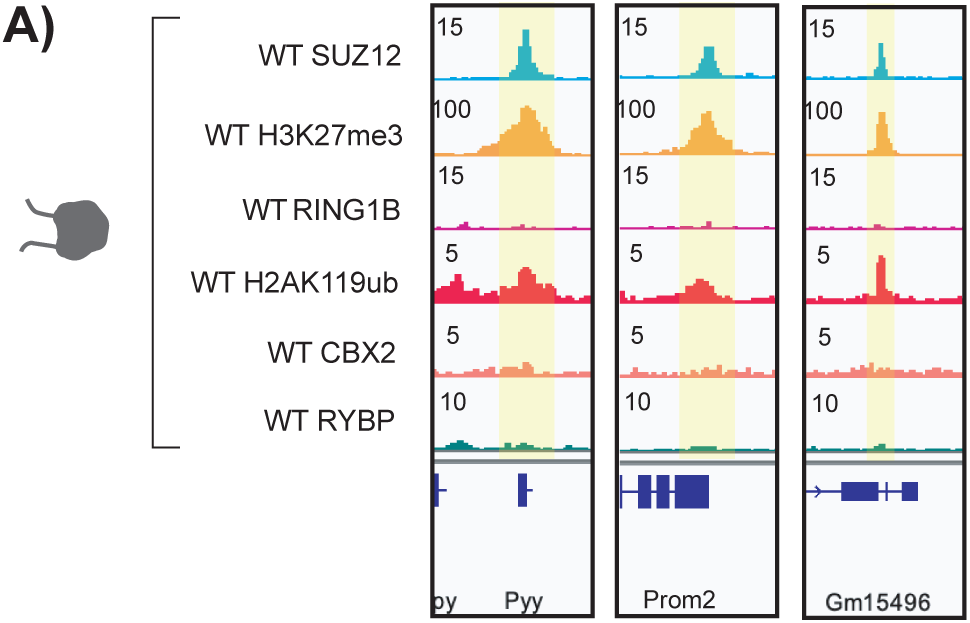

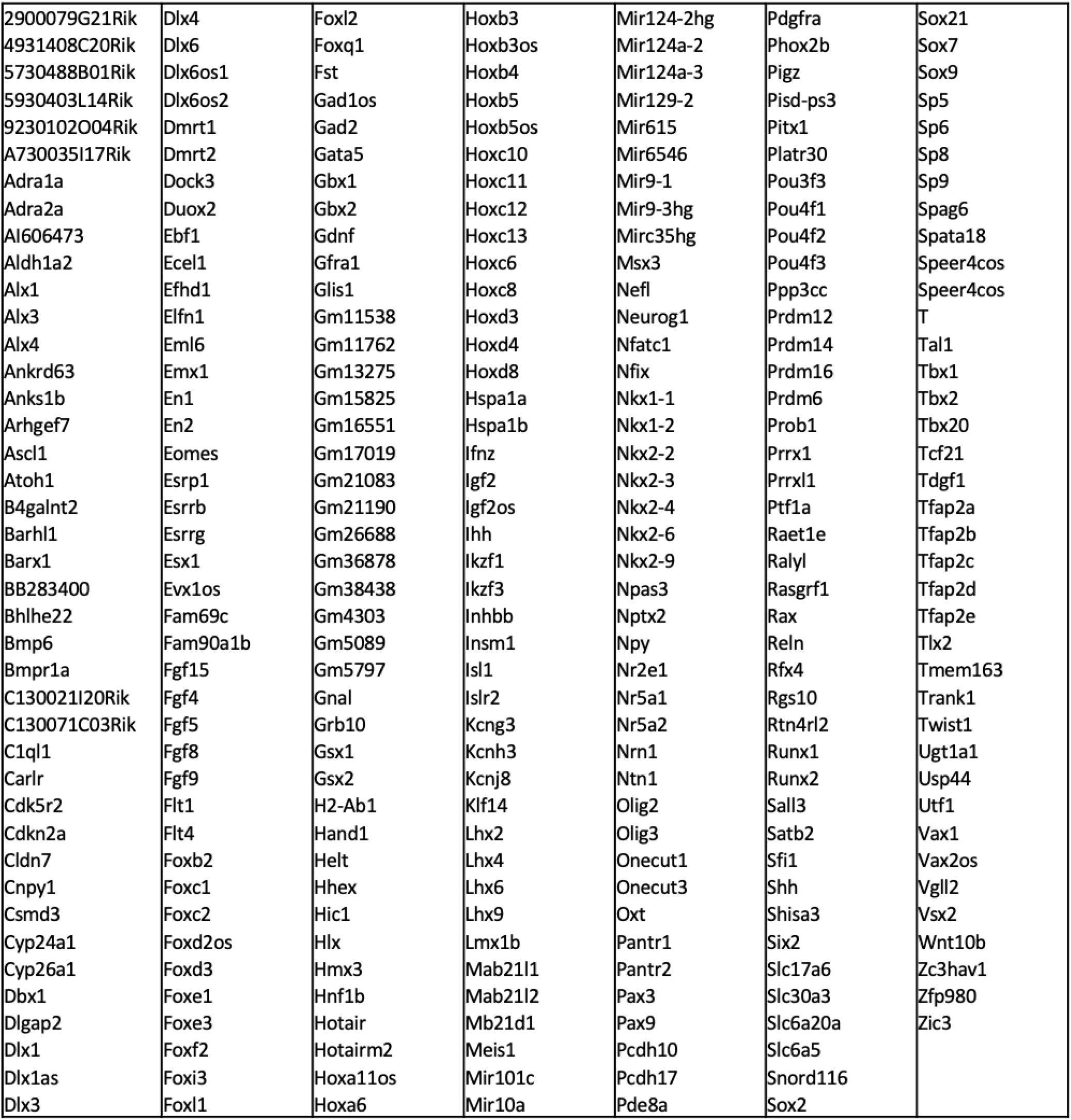

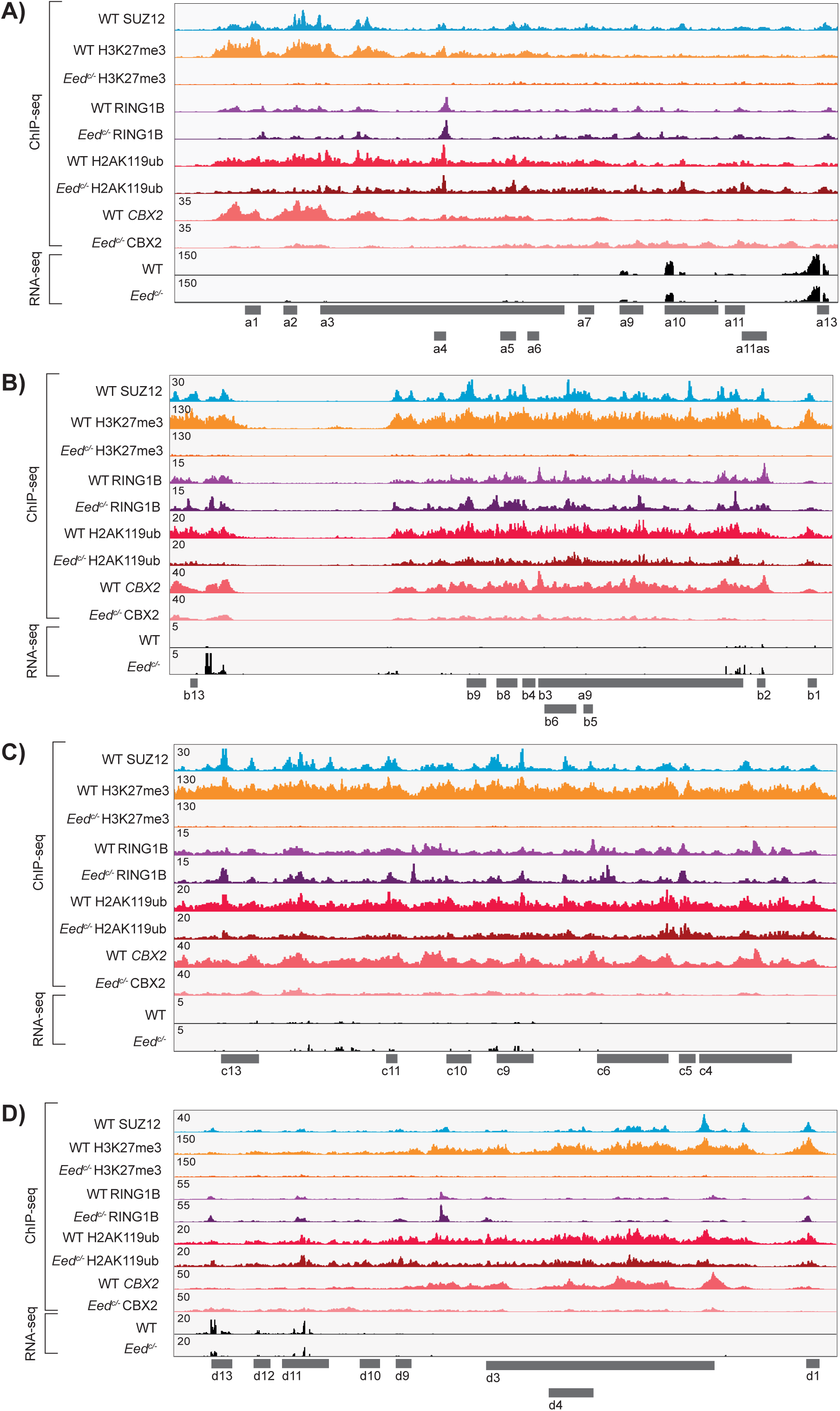

## References Table 1 and Figure 3B

Aldh1a2

[Embryonic retinoic acid synthesis is required for forelimb growth and anteroposterior patterning in the mouse](https://pubmed.ncbi.nlm.nih.gov/12117807/).

Niederreither K, **Vermot J**, Schuhbaur B, Chambon P, Dollé P. Development. 2002 Aug;129(15):3563-74.

Alx1

[Scapula development is governed by genetic interactions of Pbx1 with its family members and with Emx2 via their cooperative control of Alx1](https://pubmed.ncbi.nlm.nih.gov/20627960/).

Capellini TD, Vaccari G, Ferretti E, Fantini S, He M, Pellegrini M, Quintana L, Di Giacomo G, Sharpe J, Selleri L, Zappavigna V. Development. 2010 Aug 1;137(15):2559-69. doi: 10.1242/dev.048819.

Alx3

[BMP-2 Induced Expression of Alx3 That Is a Positive Regulator of Osteoblast Differentiation](https://pubmed.ncbi.nlm.nih.gov/23825702/).

Matsumoto T, Yamada A, Aizawa R, Suzuki D, Tsukasaki M, Suzuki W, Nakayama M, Maki K, Yamamoto M, Baba K, Kamijo R. PLoS One. 2013 Jun 18;8(6):e68774. doi: 10.1371/journal.pone.0068774. Print 2013.

Alx4

[Mutations in mouse Aristaless-like4 cause Strong’s luxoid polydactyly](https://pubmed.ncbi.nlm.nih.gov/9636085/).

Qu S, Tucker SC, Ehrlich JS, Levorse JM, Flaherty LA, Wisdom R, Vogt TF. Development. 1998 Jul;125(14):2711-21.

Atoh1

Origin of a Non-Clarke’s Column Division of the Dorsal Spinocerebellar Tract and the Role of Caudal Proprioceptive Neurons in Motor Function

<u>Rachel Yuengert ^1^, Kei Hori ^1^, Erin E Kibodeaux ^1^, Jacob X McClellan ^2^, Justin E Morales ^2^ Teng-Wei P Huang ^3^, Jeffrey L Neul ^3^, Helen C Lai ^4^</u>

Barx1

Barx1 inhibitory effect on chondrogenic initiation

Expression and function of Bapx1 during chick limb development

<u>Vicki Church ^1^, Kumiko Yamaguchi, Patricia Tsang, Keiichi Akita, Cairine Logan, Philippa Francis-West</u>

Dlx6

[The Dlx5 and Dlx6 homeobox genes are essential for craniofacial, axial, and appendicular skeletal development](https://pubmed.ncbi.nlm.nih.gov/12000792/).

Robledo RF, Rajan L, Li X, Lufkin T. Genes Dev. 2002 May 1;16(9):1089-101. doi: 10.1101/gad.988402.

Ebf1

[Expression and function of Ebf1 gene during chondrogenesis in chick embryo limb buds](https://pubmed.ncbi.nlm.nih.gov/34384862/).

El-Magd MA, Abdelfattah-Hassan A, Elsisy RA, Hawsawi YM, Oyouni AA, Al-Amer OM, El-Shetry ES. Gene. 2021 Nov 30;803:145895. doi: 10.1016/j.gene.2021.145895. Epub 2021 Aug 9.

Ecel1

[Distinct functional consequences of ECEL1/DINE missense mutations in the pathogenesis of congenital contracture disorders](https://pubmed.ncbi.nlm.nih.gov/29132416/).

Nagata K, Takahashi M, Kiryu-Seo S, Kiyama H, Saido TC. Acta Neuropathol Commun. 2017 Nov 13;5(1):83. doi: 10.1186/s40478-017-0486-9.

En1

[Multiple developmental defects in Engrailed-1 mutant mice: an early mid-hindbrain deletion and patterning defects in forelimbs and sternum](https://pubmed.ncbi.nlm.nih.gov/7925010/).

Wurst W, Auerbach AB, Joyner AL. Development. 1994 Jul;120(7):2065-75.

En2

[Examining pattern formation in mouse, chicken and frog embryos with an En-specific antiserum](https://pubmed.ncbi.nlm.nih.gov/1680044/).

Davis CA, Holmyard DP, Millen KJ, Joyner AL. Development. 1991 Feb;111(2):287-98.

Fgf4

[FGF-4 and BMP-2 have opposite effects on limb growth](https://pubmed.ncbi.nlm.nih.gov/8421496/).

Niswander L, Martin GR. Nature. 1993 Jan 7;361(6407):68-71. doi: 10.1038/361068a0.

Fgf5

[FGF5 stimulates expansion of connective tissue fibroblasts and inhibits skeletal muscle development in the limb](https://pubmed.ncbi.nlm.nih.gov/11066093/).

Clase KL, Mitchell PJ, Ward PJ, Dorman CM, Johnson SE, Hannon K. Dev Dyn. 2000 Nov;219(3):368-80. doi: 10.1002/1097-0177(2000)9999:9999<::AID-DVDY1056>3.0.CO;2-8.

Fgf8

[A role for FGF-8 in the initiation and maintenance of vertebrate limb bud outgrowth](https://pubmed.ncbi.nlm.nih.gov/7583127/).

Mahmood R, Bresnick J, Hornbruch A, Mahony C, Morton N, Colquhoun K, Martin P, Lumsden A, Dickson C, Mason I. Curr Biol. 1995 Jul 1;5(7):797-806. doi: 10.1016/s0960-9822(95)00157-6.

Fgf9

[FGF9 regulates early hypertrophic chondrocyte differentiation and skeletal vascularization in the developing stylopod](https://pubmed.ncbi.nlm.nih.gov/17544391/).

Hung IH, Yu K, Lavine KJ, Ornitz DM. Dev Biol. 2007 Jul 15;307(2):300-13. doi: 10.1016/j.ydbio.2007.04.048. Epub 2007 May 6.

Flt1

The forming limb skeleton serves as a signaling center for limb vasculature patterning via regulation of Vegf

<u>Idit Eshkar-Oren ^1^, Sergey V Viukov, Sharbel Salameh, Sharon Krief, Chun-do Oh, Haruhiko Akiyama, Hans-Peter Gerber, Napoleone Ferrara, Elazar Zelzer</u>

Foxc1

Brachydactyly type E in an Italian family with 6p25 trisomy

<u>Paolo Fontana ^1^, Cristina Tortora ^2^, Roberta Petillo ^3^, Michela Malacarne ^4^, Simona Cavani ^4^ Martina Miniero ^2^, Paola D’Ambrosio ^3^, Davide De Brasi ^5^, Maria Antonietta Pisanti ^5^</u>

Gdnf

[Neuroprotective effects of glial cell line-derived neurotrophic factor mediated by an adeno-associated virus vector in a transgenic animal model of amyotrophic lateral sclerosis](https://pubmed.ncbi.nlm.nih.gov/12177190/).

Wang LJ, Lu YY, Muramatsu S, Ikeguchi K, Fujimoto K, Okada T, Mizukami H, Matsushita T, Hanazono Y, Kume A, Nagatsu T, Ozawa K, Nakano I. J Neurosci. 2002 Aug 15;22(16):6920-8. doi: 10.1523/JNEUROSCI.22-16-06920.2002.

Hand1

[Defective Hand1 phosphoregulation uncovers essential roles for Hand1 in limb morphogenesis](https://pubmed.ncbi.nlm.nih.gov/28576769/).

Firulli BA, Milliar H, Toolan KP, Harkin J, Fuchs RK, Robling AG, Firulli AB. Development.

2017 Jul 1;144(13):2480-2489. doi: 10.1242/dev.149963. Epub 2017 Jun 2

Hic1

[Mice deficient in the candidate tumor suppressor gene Hic1 exhibit developmental defects of structures affected in the Miller-Dieker syndrome](https://pubmed.ncbi.nlm.nih.gov/10655551/).

Carter MG, Johns MA, Zeng X, Zhou L, Zink MC, Mankowski JL, Donovan DM, Baylin SB. Hum Mol Genet. 2000 Feb 12;9(3):413-9. doi: 10.1093/hmg/9.3.413.

Hoxa11as

[Evolution of Hoxa11 regulation in vertebrates is linked to the pentadactyl state](https://pubmed.ncbi.nlm.nih.gov/27706137/).

Kherdjemil Y, Lalonde RL, Sheth R, Dumouchel A, de Martino G, Pineault KM, Wellik DM, Stadler HS, Akimenko MA, Kmita M. Nature. 2016 Nov 3;539(7627):89-92. doi: 10.1038/nature19813. Epub 2016 Oct 5.

Ihh

[Regulation of rate of cartilage differentiation by Indian hedgehog and PTH-related protein](https://pubmed.ncbi.nlm.nih.gov/8662546/).

Vortkamp A, Lee K, Lanske B, Segre GV, Kronenberg HM, Tabin CJ. Science. 1996 Aug 2;273(5275):613-22. doi: 10.1126/science.273.5275.613.

Lhx2

[LIM homeobox transcription factors integrate signaling events that control three-dimensional limb patterning and growth](https://pubmed.ncbi.nlm.nih.gov/19304889/).

Tzchori I, Day TF, Carolan PJ, Zhao Y, Wassif CA, Li L, Lewandoski M, Gorivodsky M, Love PE, Porter FD, Westphal H, Yang Y. Development. 2009 Apr;136(8):1375-85. doi: 10.1242/dev.026476.

Lmx1b

[Mutations in LMX1B cause abnormal skeletal patterning and renal dysplasia in nail patella syndrome](https://pubmed.ncbi.nlm.nih.gov/9590287/).

Dreyer SD, Zhou G, Baldini A, Winterpacht A, Zabel B, Cole W, Johnson RL, Lee B. Nat Genet. 1998 May;19(1):47-50. doi: 10.1038/ng0598-47.

Meis1

[Conserved regulation of proximodistal limb axis development by Meis1/Hth](https://pubmed.ncbi.nlm.nih.gov/10586884/).

Mercader N, Leonardo E, Azpiazu N, Serrano A, Morata G, Martínez C, Torres M. Nature. 1999 Nov 25;402(6760):425-9. doi: 10.1038/46580.

Pax3

[Pax-3 is required for the development of limb muscles: a possible role for the migration of dermomyotomal muscle progenitor cells](https://pubmed.ncbi.nlm.nih.gov/8162858/).

Bober E, Franz T, Arnold HH, Gruss P, Tremblay P. Development. 1994 Mar;120(3):603-12.

Lhx9

[LIM homeobox transcription factors integrate signaling events that control three-dimensional limb patterning and growth](https://pubmed.ncbi.nlm.nih.gov/19304889/).

Tzchori I, Day TF, Carolan PJ, Zhao Y, Wassif CA, Li L, Lewandoski M, Gorivodsky M, Love PE, Porter FD, Westphal H, Yang Y.

Pax9

[Pax9-deficient mice lack pharyngeal pouch derivatives and teeth and exhibit craniofacial and limb abnormalities](https://pubmed.ncbi.nlm.nih.gov/9732271/).

Peters H, Neubüser A, Kratochwil K, Balling R. Genes Dev. 1998 Sep 1;12(17):2735-47. doi: 10.1101/gad.12.17.2735.

Pitx1

[Role of the Bicoid-related homeodomain factor Pitx1 in specifying hindlimb morphogenesis and pituitary development](https://pubmed.ncbi.nlm.nih.gov/10049363/).

Szeto DP, Rodriguez-Esteban C, Ryan AK, O’Connell SM, Liu F, Kioussi C, Gleiberman AS, Izpisúa-Belmonte JC, Rosenfeld MG. Genes Dev. 1999 Feb 15;13(4):484-94. doi: 10.1101/gad.13.4.484.

Prx1

[Prx1 and Prx2 in skeletogenesis: roles in the craniofacial region, inner ear and limbs](https://pubmed.ncbi.nlm.nih.gov/9729491/).

ten Berge D, Brouwer A, Korving J, Martin JF, Meijlink F. Development. 1998 Oct;125(19):3831-42.

Ptf1a

[Genome-wide chromatin accessibility and transcriptome profiling show minimal epigenome changes and coordinated transcriptional dysregulation of hedgehog signaling in Danforth’s short tail mice](https://pubmed.ncbi.nlm.nih.gov/30380057/).

Orchard P, White JS, Thomas PE, Mychalowych A, Kiseleva A, Hensley J, Allen B, Parker SCJ, Keegan CE. Hum Mol Genet. 2019 Mar 1;28(5):736-750. doi: 10.1093/hmg/ddy378.

Runx1

[Runx1/AML1/Cbfa2 mediates onset of mesenchymal cell differentiation toward chondrogenesis](https://pubmed.ncbi.nlm.nih.gov/16059634/).

Wang Y, Belflower RM, Dong YF, Schwarz EM, O’Keefe RJ, Drissi H. J Bone Miner Res. 2005 Sep;20(9):1624-36. doi: 10.1359/JBMR.050516. Epub 2005 May 23.

Runx2

[Runx2 and Runx3 are essential for chondrocyte maturation, and Runx2 regulates limb growth through induction of Indian hedgehog](https://pubmed.ncbi.nlm.nih.gov/15107406/).

Yoshida CA, Yamamoto H, Fujita T, Furuichi T, Ito K, Inoue K, Yamana K, Zanma A, Takada K, Ito Y, Komori T. Genes Dev. 2004 Apr 15;18(8):952-63. doi: 10.1101/gad.1174704.

Sall3

[Sall genes regulate region-specific morphogenesis in the mouse limb by modulating Hox activities](https://pubmed.ncbi.nlm.nih.gov/19168674/).

Kawakami Y, Uchiyama Y, Rodriguez Esteban C, Inenaga T, Koyano-Nakagawa N, Kawakami H, Marti M, Kmita M, Monaghan-Nichols P, Nishinakamura R, Izpisua Belmonte JC. Development. 2009 Feb;136(4):585-94. doi: 10.1242/dev.027748.

Shh

[A positive feedback loop coordinates growth and patterning in the vertebrate limb](https://pubmed.ncbi.nlm.nih.gov/7935794/).

Niswander L, Jeffrey S, Martin GR, Tickle C. Nature. 1994 Oct 13;371(6498):609-12. doi: 10.1038/371609a0.

Six2

[Wnt and BMP signaling cooperate with Hox in the control of Six2 expression in limb tendon precursor](https://pubmed.ncbi.nlm.nih.gov/23499659/).

Yamamoto-Shiraishi Y, Kuroiwa A. Dev Biol. 2013 May 15;377(2):363-74. doi: 10.1016/j.ydbio.2013.02.023. Epub 2013 Mar 13.

Sp6

[The development of several organs and appendages is impaired in mice lacking Sp6](https://pubmed.ncbi.nlm.nih.gov/18297738/).

Hertveldt V, Louryan S, van Reeth T, Drèze P, van Vooren P, Szpirer J, Szpirer C. Dev Dyn. 2008 Apr;237(4):883-92. doi: 10.1002/dvdy.21355.

Sp8

[Sp8 is crucial for limb outgrowth and neuropore closure](https://pubmed.ncbi.nlm.nih.gov/14526104/).

Bell SM, Schreiner CM, Waclaw RR, Campbell K, Potter SS, Scott WJ. Proc Natl Acad Sci U S A. 2003 Oct 14;100(21):12195-200. doi: 10.1073/pnas.2134310100. Epub 2003 Oct 2.

Sp9

[Sp8 and Sp9, two closely related buttonhead-like transcription factors, regulate Fgf8 expression and limb outgrowth in vertebrate embryos](https://pubmed.ncbi.nlm.nih.gov/15358670/).

Kawakami Y, Esteban CR, Matsui T, Rodríguez-León J, Kato S, Izpisúa Belmonte JC. Development. 2004 Oct;131(19):4763-74. doi: 10.1242/dev.01331.

Tbx1

[Tbx1 regulation of myogenic differentiation in the limb and cranial mesoderm](https://pubmed.ncbi.nlm.nih.gov/17117436/).

Dastjerdi A, Robson L, Walker R, Hadley J, Zhang Z, Rodriguez-Niedenführ M, Ataliotis P, Baldini A, Scambler P, Francis-West P. Dev Dyn. 2007 Feb;236(2):353-63. doi: 10.1002/dvdy.21010.

Tfap2a

[Search for genetic modifiers of IRF6 and genotype-phenotype correlations in Van der Woude and popliteal pterygium syndromes](https://pubmed.ncbi.nlm.nih.gov/23949966/).

Leslie EJ, Mancuso JL, Schutte BC, Cooper ME, Durda KM, L’Heureux J, Zucchero TM, Marazita ML, Murray JC. Am J Med Genet A. 2013 Oct;161A(10):2535-2544. doi: 10.1002/ajmg.a.36133.

Epub 2013 Aug 15.

Tfap2b

[Mutations in TFAP2B cause Char syndrome, a familial form of patent ductus arteriosus](https://pubmed.ncbi.nlm.nih.gov/10802654/).

Satoda M, Zhao F, Diaz GA, Burn J, Goodship J, Davidson HR, Pierpont ME, Gelb BD. Nat Genet. 2000 May;25(1):42-6. doi: 10.1038/75578.

Tfap2c (suspected role)

[A discrete transition zone organizes the topological and regulatory autonomy of the adjacent tfap2c and bmp7 genes](https://pubmed.ncbi.nlm.nih.gov/25569170/).

Tsujimura T, Klein FA, Langenfeld K, Glaser J, Huber W, Spitz F. PLoS Genet. 2015 Jan 8;11(1):e1004897. doi: 10.1371/journal.pgen.1004897. eCollection 2015 Jan.

Twist1

[Unraveling the transcriptional regulation of TWIST1 in limb development](https://pubmed.ncbi.nlm.nih.gov/30372441/).

Hirsch N, Eshel R, Bar Yaacov R, Shahar T, Shmulevich F, Dahan I, Levaot N, Kaplan T, Lupiáñez DG, Birnbaum RY. PLoS Genet. 2018 Oct 29;14(10):e1007738. doi: 10.1371/journal.pgen.1007738.

Wnt10b

[Homozygous WNT10b mutation and complex inheritance in Split-Hand/Foot Malformation](https://pubmed.ncbi.nlm.nih.gov/18515319/).

Ugur SA, Tolun A. Hum Mol Genet. 2008 Sep 1;17(17):2644-53. doi: 10.1093/hmg/ddn164. Epub 2008 May 30.

Zic3

[Preaxial polydactyly caused by Gli3 haploinsufficiency is rescued by Zic3 loss of function in mice](https://pubmed.ncbi.nlm.nih.gov/22234993/).

Quinn ME, Haaning A, Ware SM. Hum Mol Genet. 2012 Apr 15;21(8):1888-96. doi: 10.1093/hmg/dds002. Epub 2012 Jan 10.

Bmpr1a

[BMP signals control limb bud interdigital programmed cell death by regulating FGF signaling](https://pubmed.ncbi.nlm.nih.gov/17537800/).

Pajni-Underwood S, Wilson CP, Elder C, Mishina Y, Lewandoski M. Development. 2007 Jun;134(12):2359-68. doi: 10.1242/dev.001677.

Cyp26b1

[Regulation of retinoic acid distribution is required for proximodistal patterning and outgrowth of the developing mouse limb](https://pubmed.ncbi.nlm.nih.gov/15030763/).

Yashiro K, Zhao X, Uehara M, Yamashita K, Nishijima M, Nishino J, Saijoh Y, Sakai Y, Hamada H. Dev Cell. 2004 Mar;6(3):411-22. doi: 10.1016/s1534-5807(04)00062-0.

Hoxa9, a10, a11, a13 See references in

[The role of Hox genes during vertebrate limb development](https://pubmed.ncbi.nlm.nih.gov/17644373/).

**Zakany J**, **Duboule D.** Curr Opin Genet Dev. 2007 Aug;17(4):359-66. doi: 10.1016/j.gde.2007.05.011. Epub 2007 Jul 20.

Mecom

[Lethal neonatal bone marrow failure syndrome with multiple congenital abnormalities, including limb defects, due to a constitutional deletion of 3’ MECOM](https://pubmed.ncbi.nlm.nih.gov/29439187/).

van der Veken LT, Maiburg MC, Groenendaal F, van Gijn ME, Bloem AC, Erpelinck C, Gröschel S, Sanders MA, Delwel R, Bierings MB, Buijs A. Haematologica. 2018 Apr;103(4):e173-e176. doi: 10.3324/haematol.2017.185033. Epub 2018 Feb 8.

Pdgfra

[Acquisition of a Unique Mesenchymal Precursor-like Blastema State Underlies Successful Adult Mammalian Digit Tip Regeneration](https://pubmed.ncbi.nlm.nih.gov/31902657/).

Storer MA, Mahmud N, Karamboulas K, Borrett MJ, Yuzwa SA, Gont A, Androschuk A, Sefton MV, Kaplan DR, Miller FD. Dev Cell. 2020 Feb 24;52(4):509-524.e9. doi: 10.1016/j.devcel.2019.12.004. Epub 2020 Jan 2.

Tbx5

[Tbx5 is required for forelimb bud formation and continued outgrowth](https://pubmed.ncbi.nlm.nih.gov/12736217/).

Rallis C, Bruneau BG, Del Buono J, Seidman CE, Seidman JG, Nissim S, Tabin CJ, Logan MP. Development. 2003 Jun;130(12):2741-51. doi: 10.1242/dev.00473.

Yy1

[YY1 activates Msx2 gene independent of bone morphogenetic protein signaling](https://pubmed.ncbi.nlm.nih.gov/11861914/).

Tan DP, Nonaka K, Nuckolls GH, Liu YH, Maxson RE, Slavkin HC, Shum L. Nucleic Acids Res. 2002 Mar 1;30(5):1213-23. doi: 10.1093/nar/30.5.1213.

## Notes

### Competing Interest Statement

The authors have declared no competing interest.

